# Consistent Under-reporting of Task Details in Motor Imagery Research

**DOI:** 10.1101/2022.10.25.513501

**Authors:** Elise E Van Caenegem, Gautier Hamoline, Baptiste M Waltzing, Robert M Hardwick

## Abstract

Motor Imagery is a subject of longstanding scientific interest. However, critical details of motor imagery protocols are not always reported in full, hampering direct replication and translation of this work. The present review provides a quantitative assessment of the prevalence of under-reporting in the recent motor imagery literature. Publications from the years 2018-2020 were examined, with 695 meeting the inclusion criteria for further examination. Of these studies, 64% (445/695) did not provide information about the modality of motor imagery (i.e., kinesthetic, visual, or a mixture of both) used in the study. When visual or mixed imagery was specified, the details of the visual perspective to be used (i.e., first person, third person, or combinations of both) were not reported in 24% (25/103) of studies. Further analysis indicated that studies using questionnaires to assess motor imagery reported more information than those that did not. We conclude that studies using motor imagery consistently under-report key details of their protocols, which poses a significant problem for understanding, replicating, and translating motor imagery effects.

## 1. Introduction

The effects of motor imagery (i.e., imagining the execution of an action without physically performing it) have been studied for many years (e.g., James, 1890). Motor imagery can be used to increase athletic performance (Ladda et al., 2021), to learn new skills (Lotze & Halsband, 2006; Williams & Gribble, 2012), for rehabilitation (Malouin & Richards, 2010; Jackson et al., 2001; Ietswaart et al., 2011), in brain computer interfaces studies (Chaudhary et al., 2016), and many other domains. As such, there is long-lasting and multidisciplinary scientific interest in motor imagery.

‘Motor Imagery’ can itself be considered a blanket term; there are many different ways in which a movement can be imagined, with the kinesthetic and visual modalities being the most relevant to motor imagery in the scientific literature (McAvinue & Robertson, 2008). Kinesthetic imagery can be defined as a ‘representation of the sensations experienced during physical performance including muscle tension, proprioception, force and effort involved in movement’ (Callow & Waters, 2005), and often involves instructions that emphasise ‘feeling’ the movement, or focusing on the sensations that the movement generates. By contrast, visual imagery has been defined as ‘the representation of perceptual information in the absence of visual input’ (Kaski, 2002) which can include visualisation of body movements and aspects of the external environment. Visual imagery can be further characterised by adopting a first-person perspective, in which the movement is imaged as if the individual were seeing through their own eyes, or a third person perspective, in which the movement is visualized from outside the body. These different forms of imagery are not directly equivalent and have dissociable neural effects. Comparisons of kinesthetic and visual imagery have found that they respectively recruit regions more closely associated with motor functions and visual perception (Guillot et al., 2009), and that kinesthetic imagery modulates corticospinal excitability, while visual motor imagery does not (Hardy & Callow, 1999; Jiang et al., 2015; Kilintari et al., 2016; Lee et al., 2019; Seiler et al., 2015; Stinear et al., 2006). Similarly, Jiang et al. (2015) demonstrated differentiated neuronal activation for the different perspectives, with internal imagery (which combines first person and kinesthetic perspectives) recruiting more brain regions than third person imagery. As such, in order to accurately communicate the protocol used in research using motor imagery, it is important for researchers to clarify details such as the modality (kinesthetic, visual, or a mixture of both) and visual perspective (first person, third person, or a mixture of both) from which actions are imagined.

Recent work has highlighted the issue of under-reporting of protocol details in the motor imagery literature. For example, a meta-analysis of neuroimaging studies found that that approximately 2/3 of papers did not provide enough information to identify the visual perspective that participants used during motor imagery (Hardwick et al., 2018). Studies attempting to complete systematic reviews of the literature have also noted that critical details allowing the full understanding of the protocols used are lacking (Silva et al., 2020, Baniqued et al. 2021). However, the central goal of these studies was not to examine the propensity of under-reporting in the motor imagery literature; as such, the extent and prevalence of this issue remains unclear. The goal of the present study was therefore to conduct a systematic analysis to examine the issue of underreporting in recent motor imagery publications. We examined two central questions; first, whether studies reported all information required to identify the modality of imagery used (kinesthetic, visual or both), and second, whether those studies that included visual motor imagery included enough information to identify the perspective used (1^st^ person, 3^rd^ person or both). Finally, as questionnaires (McAvinue & Robertson, 2008, Malouin et al., 2007, Isaac et al., 1986) that assess motor imagery ability often prompt users to consider the modality and perspective of the imagery used (e.g., KVIQ), we asked whether those studies that included a questionnaire reported more information than those that did not.

## 2. Methods

### 2.1 Literature searches

Papers were found through PubMed literature searches. A search for papers on Motor Imagery was conducted using the search string ‘mental imagery’ OR ‘kinesthetic imagery’ OR ‘motor imagery’ OR ‘visual imagery’ OR ‘mental practice’ OR ‘mental training’ OR ‘mental rehearsal’. This research string was based on the terms used in a previous study by Guerra et al. (2017). The search was limited to the years 2018, 2019 and 2020 to enable the feasibility of the study. These criteria provided a total of 1700 papers (n=530 for the year 2018, n=540 for 2019, and n=630 for 2020.

### 2.2 Inclusion/Exclusion criteria

By reading the abstracts, studies that did not directly involve motor imagery (e.g., reviews and meta-analyses) were removed. Duplicates and articles not available in English were also removed. We also removed papers studying implicit motor imagery (because these protocols do not provide motor imagery instructions to participants) and studies which involved action observation coupled with motor imagery, i.e., ‘AOMI’ (Vogt et al., 2013) (as they provide a visual stimulus and so emphasize kinesthetic imagery, making classification unnecessary). Of the 1700 papers screened, 695 articles met the criteria for further analysis (2018 n=219, 2019 n=209, 2020 n=267) (Fig. 1).

**Figure 1:**
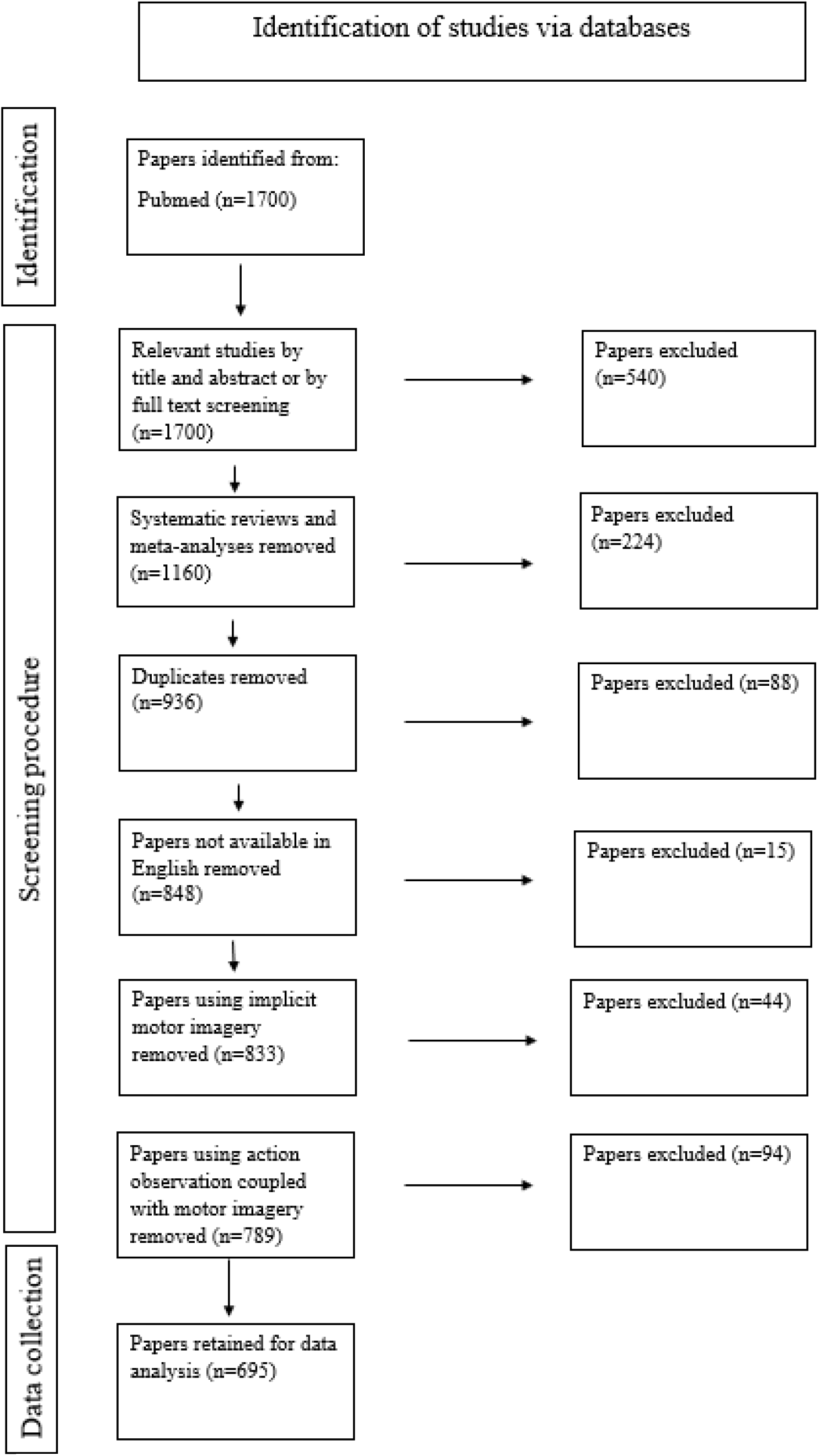
Screening procedure

### 2.3 Data encoding

While screening papers, we extracted data that allowed us to answer three questions. Our first and central question regarded whether the publication included enough information to allow us to determine the modality of motor imagery used. Papers were classified according to whether the modality was identifiable as ‘kinesthetic’, ‘visual’, ‘mixed’ (i.e., kinesthetic, and visual), or ‘not stated’. Our second question examined whether papers that included the use of visual motor imagery (i.e., those encoded as using a modality that was ‘visual’ or ‘mixed’) also reported enough information to determine the visual perspective that was used. Data were encoded as ‘first person’, ‘third person’, ‘mixed’ (e.g., when both first and third person perspectives were used in the same experiment), or ‘not stated’. Our third and final question examined whether studies that included questionnaires to assess motor imagery ability were also more likely to report details of the modality and/or perspective of motor imagery used in their protocol. These data included the name of the questionnaire (e.g., MIQ, etc), or ‘none’ if no questionnaire was used. Data extracted, including links to each paper, are reported in the supplementary materials for this publication (see supplementary materials).

### 2.4 Statistical analysis

Statistical analyses were conducted using JASP Version 0.16 (JASP Team 2021). We first examined the reporting of the modality of motor imagery; a binomial test examined whether the proportion of studies for which the modality of imagery could be identified was lower than 1.0 (representing the ideal that all studies should report this information). Similarly, a second analysis used a binomial test to determine whether studies using visual or mixed modalities of motor imagery also included information about the visual perspective used, examining whether the proportion of studies in which the visual perspective could be identified was lower than 1.0. Finally, chi-squared tests examined whether the inclusion of a questionnaire affected the likelihood that these details were reported.

Each analysis was examined using two forms of statistical test. First, an analysis was conducted using a classical frequentist statistical test, with an alpha value of p<0.05. As frequentist tests represent the most commonly used form of statistical analysis, this allows the presentation of the results in a widely understood and accessible format. However, frequentist statistics face a limitation that is particularly relevant to the interpretation of the present study; they can be used only to reject the null hypothesis and cannot be used to provide evidence that the null hypothesis is true. As such, if our analysis found that under-reporting was actually a relatively minimal problem in the motor imagery literature, the results of frequentist tests would be inconclusive. By contrast, Bayesian analyses can be used to actively quantify how much more likely one of these hypotheses is than the other. The Bayes Factor (BF) presents a ratio of evidence for the alternative hypothesis (BF_10_) in comparison to the null hypothesis (BF_01_), with values of BF_10_>1 representing evidence in favour of the alternative hypothesis. This had led researchers to consider the use of a certain Bayes Factor as an appropriate ‘stopping criterion’ (typically BF_10_>30, representing ‘strong evidence’ in favour of the alternative hypothesis), at which it can be concluded that evidence is compelling enough to support the hypothesis without further need for data collection (Wagenmakers et al., 2019). As such, we conducted additional Bayesian tests to determine whether the data from our present sample, which was limited to the years 2018-2020, would allow us to provide a conclusive answer to our research questions.

## 3. Results

### 3.1 Reporting of Imagery Modality

Our first analysis examined whether there was evidence of under-reporting of the details required to understand the modality of imagery used (Kinesthetic, Visual, Mixed) in the published literature. A binomial test was used to examine this question, with the aim to compare the proportions observed in our sample with the ‘null’ hypothesis that all papers did provide adequate detail (i.e., the proportion ‘Yes’ = 1.00). The analysis indicated that a significant majority of the papers did not provide sufficient information to determine the modality of imagery that was used in a way that would allow full understanding or future replication of the procedures used (445/695, 64%, p<0.001) (Fig. 2A). Further Bayesian analysis (BF_10_=5.017e + 10) indicated the result represents ‘extreme evidence’ (Quintana & Williams, 2018; Wagenmakers et al., 2019) for the hypothesis that papers do not report adequate details. The extreme value identified here also suggests that our sample size provides sufficient evidence to conclude that most motor imagery studies do not provide adequate details of the modality of imagery used in their study protocol.

**Figure 2:**
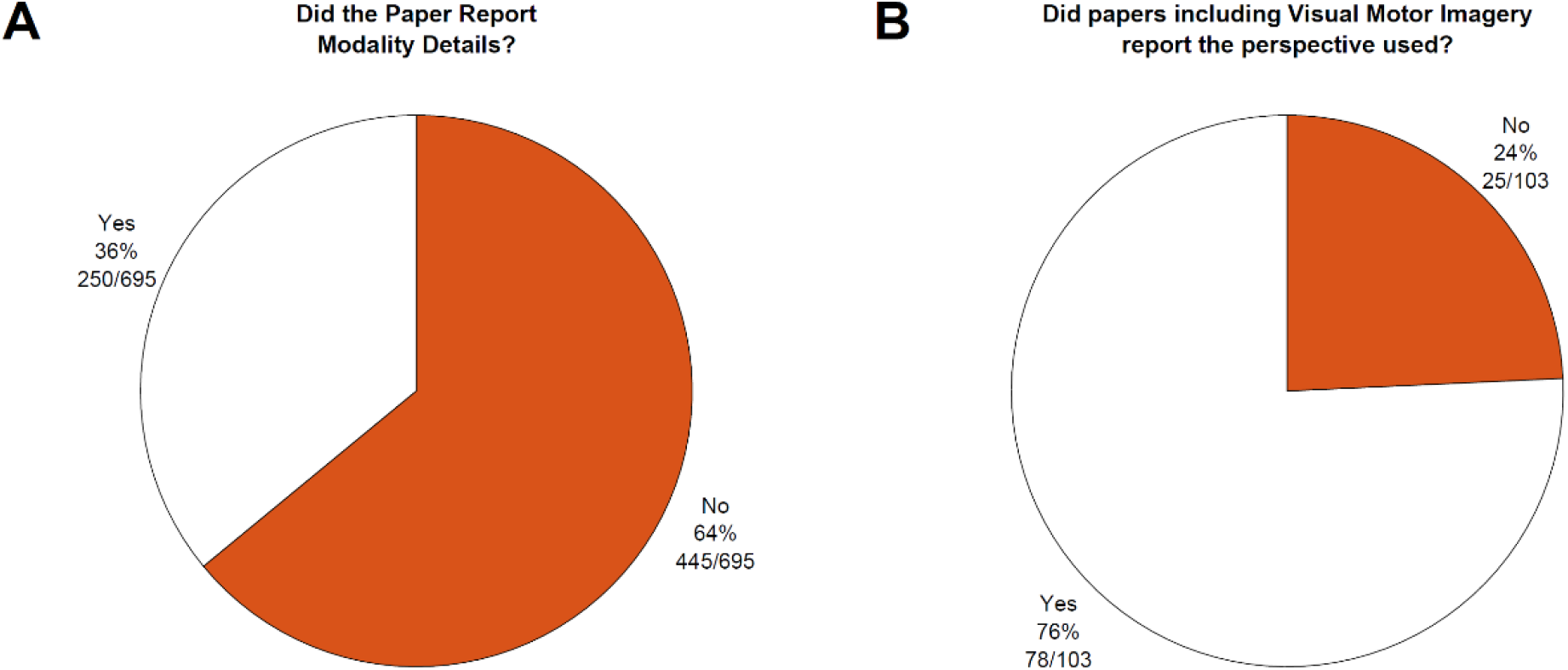
Reporting of Information in Motor Imagery Studies. A) The proportion of papers reporting full details of motor imagery. B) The proportion of papers providing perspective when visual imagery is specified.

### 3.2 Reporting of Perspective in Visual Imagery

Our second analysis examined whether articles that used a purely visual or a mixture of visual and kinesthetic motor imagery modality provided information on the visual perspective used (First person, Third person, or a mixture of both). While a majority of papers (75.7%, 78/103) did report this information, approximately one in four (24.3%, 25/103) did not, which was again consistent with our hypothesis that there is significant under-reporting in the motor imagery literature (Binomial test, p<0.001) (Fig. 2B). Further Bayesian analysis provided additional support for this conclusion (BF_10_=172,962), which represents ‘extreme evidence’ (Quintana & Williams, 2018; Wagenmakers et al., 2019) for our hypothesis that papers do not report adequate details.

### 3.3 Impact of the use of questionnaires

A chi-square test identified a significant interaction between whether studies included the use of a questionnaire to assess motor imagery ability (yes or no), and whether they reported all relevant details regarding imagery modality and perspective (yes or no); *x*^2^ =179.11, p<0.001. Further Bayesian analysis indicated that this represented ‘extreme evidence’ that the use of questionnaires led to differences in study reporting (BF_10_=4.768e+37). Studies that used a questionnaire (representing 134/695 or 19.3% of the total sample) were significantly more likely to report relevant protocol details (studies using a questionnaire that reported all details; 115/134, 85.8%; studies that did not use a questionnaire that reported all details; 142/561, 25.3%) (Fig. 3A).

**Figure 3:**
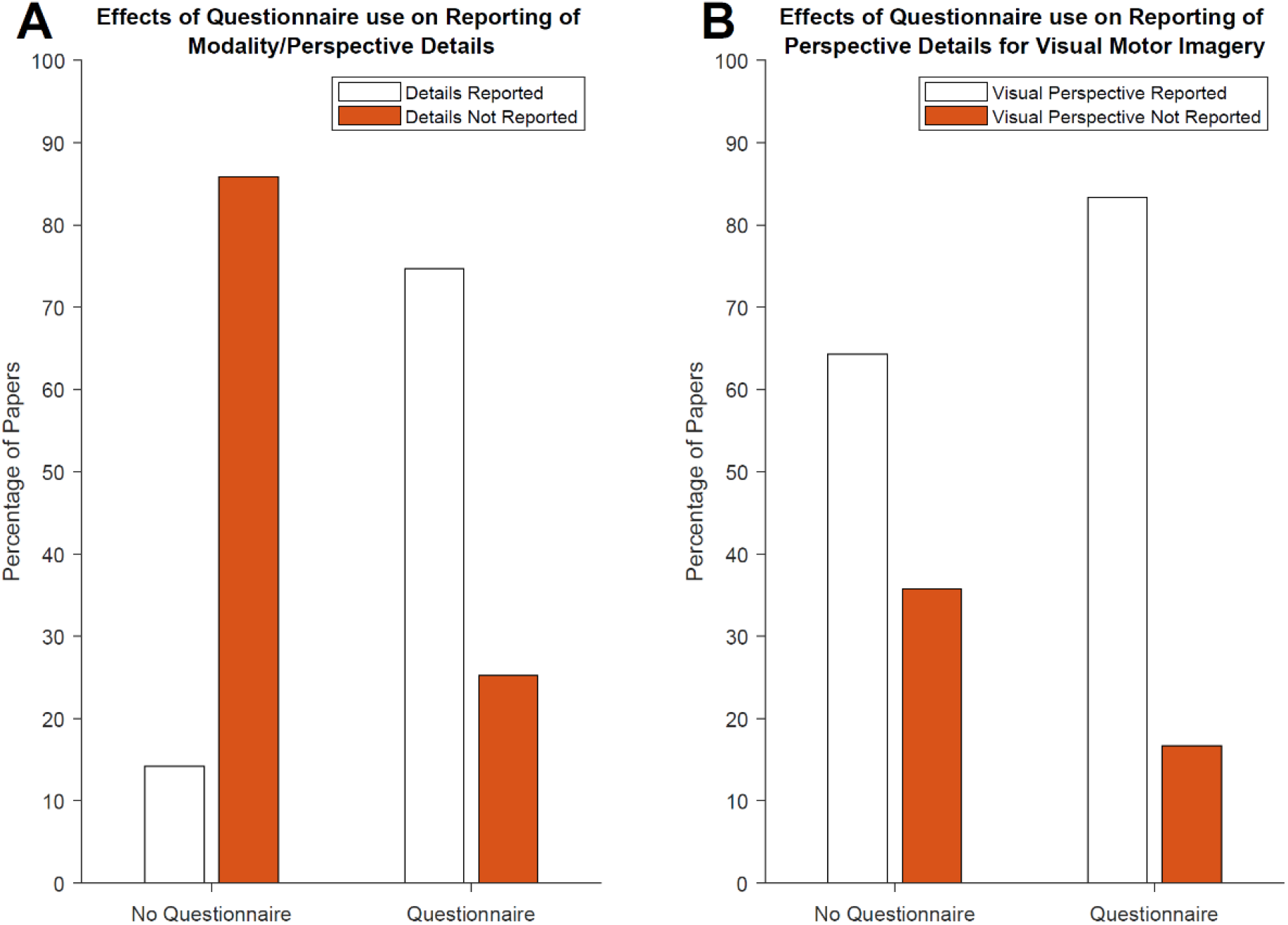
Effects of questionnaires on reporting. A) The proportion of articles reporting motor imagery details. B) The proportion of papers providing perspective when visual imagery is specified.

Further analysis examined whether the use of a questionnaire also affected the reporting of visual perspective in the 103 papers that were identified as using visual or mixed (visual and kinesthetic) motor imagery (Fig 3B). A chi-squared test identified a trend towards significance *x*^2^ = 2.739, p=0.098, BF_10_=0.889, where papers that did include a questionnaire were more likely to report the relevant details (papers that used a questionnaire and did report the visual perspective used; 60/75, 80.0%; papers that did not use a questionnaire and did report the visual perspective used; 18/28, 64.3%).

## 4. Discussion

### 4.1 Evidence of under-reporting in the motor imagery literature

The present study examined the hypothesis that articles in the recent literature on motor imagery did not provide enough information to allow full understanding of their procedures. The results of our analyses support this hypothesis. Regarding motor imagery modality, 64% of articles did not provide information on whether the study used visual, kinesthetic, or a mixture of visual and kinesthetic imagery. When papers specified that their tasks involved visual imagery, 24% articles did not provide information regarding perspective (first-person, third person or a mixture of these perspectives). These results highlight significant under-reporting of details critical to understand the procedures used in the recent motor-imagery literature.

### 4.2 Impact of the use of a questionnaire

Using a questionnaire to assess motor imagery ability seems to have a positive impact on the reporting of information in the published literature. Indeed, in 85% of the cases where a questionnaire was used, it was possible to identify the modality of the imagery used for the study (visual, kinesthetic or both), and there was also a trend whereby studies that reported using visual motor imagery were more likely to report the perspective used.

We interpret these findings as being a result of questionnaires ‘prompting’ experimenters to consider relevant details of their motor imagery protocols in greater detail. Most of the questionnaires used in these studies (MIQ, KVIQ, VMIQ) ask participants to imagine different items according to the different modalities and perspectives of motor imagery. Experimenters administrating these questionnaires are therefore more likely to be aware of the possible differences in modality and perspective that can be used during motor imagery, making them more likely to consider them when developing their study protocols and writing corresponding reports. Indeed, in many cases, what was asked to be imaged in the questionnaire was identical to what was asked in the task for which the research was being conducted. We note, however, that the use of a questionnaire did not always ensure that all relevant details relating to the motor imagery modality and perspective were reported. While the use of such questionnaires is recommended in order to provide additional characterization of the study population, additional measures are recommended to ensure that all relevant details of the study are reported in full.

### 4.3 Lack of standard reporting protocol

The systematic under-reporting observed leads us to suggest that introducing clear reporting protocols would be beneficial for study reporting in the field of motor imagery. Several fields have begun to introduce standard reporting procedures to improve the inclusion of relevant details (Chipchase et al., 2012; Moseley et al., 2002; Quintana et al., 2016). The introduction of reporting protocols in the field of Motor Imagery could therefore have several advantages. Primarily, the presence of methodological details is essential for the replication and understanding of studies, and standard reporting procedures would assist with the creation of systematic reviews and meta-analysis synthesizing work in a field. Moreover, the presence of a clearly defined reporting protocol could facilitate the peer review process, as ensuring that relevant details are present in the manuscript from an early stage would reduce back-and-forth discussions about methodological issues. We therefore anticipate that the development of standardised procedures to improve the level of detail reported in action simulation studies would be extremely beneficial for future work aimed at replicating work and translating work on motor imagery into applied and clinical contexts. Such guidelines have only recently been introduced (Moreno-Verdú et al., 2022) and while at the time of writing it is not yet possible to assess the effects of these guidelines on the motor imagery literature, we anticipate that such initiatives will have a positive effect on the quality of study reporting.

### 4.4 Inconsistent terminology across studies

The terminology used to describe motor imagery interventions differs considerably between studies. A primary concern is that the term ‘motor imagery’ is not sufficient to define the imagery modality, it is not always clear whether visual imagery, kinesthetic imagery, or a combination of both is used, which requires readers to attempt to interpret or deduce these details of the procedure. For kinesthetic imagery, the precise term is not always used; in some cases, it is mentioned to ‘focus on the sensations’ or ‘feel the movement’. Concerning perspectives, there are different terms to express the same concepts. For the first-person perspective, it is possible to find the term ‘internal perspective’ or ‘egocentric perspective’. The third-person perspective is also called the ‘external’ or ‘allocentric’ perspective. Readers unfamiliar with the field of motor imagery may therefore miss or misinterpret essential information due to the use of different terminology. As such, authors may wish to consider using more standardized terminology from previously published works (McAvinue & Robertson, 2008) when describing the procedures in their studies, or clearly defining what is intended when using specific terms.

### 4.5 Indirect reporting

In several cases, the authors of published work pointed the reader to a previously published protocol but did not provide a summary of the procedures. In these cases, we classified the imagery used based on the document that was referenced. However, it is important to stress that in the future it is preferable that studies provide such information directly in order to avoid an additional step for the reader to find the information in order to be able to more easily understand the procedures undertaken.

### 4.6 Under-reporting in open-access datasets

When performing qualitative inspections of the different studies, we found that many of the studies using brain computer-interface approaches analysed data from openly available datasets (BCI Competition IV (bbci.de)). However, we found that the majority of these datasets do not provide any information about the modality and/or perspective of the imagined actions. We therefore recommend that future datasets include more information on these points, as it would drastically improve our understanding of what participants were doing in these experiments.

### 4.7 Limitations

The present research was not a full systematic review of the literature, covering the years 2018-2020 for the purposes of feasibility. However, Bayesian analyses indicated that the present results indicate extremely strong evidence for our central hypothesis that there is significant under-reporting of protocol details in the motor imagery literature. This suggests that further analysis would not change the empirical results of our study, and helps further support the idea that there is significant under-reporting in the literature.

## 5. Conclusions

In conclusion, the results presented in this paper indicate that recent studies on motor imagery do not provide enough information about the modalities (visual, kinesthetic, mixed) and perspectives (first person, third person, mixed) of the task being performed. The use of questionnaires (MIQ, KVIQ, VMIQ, etc.) to assess motor imagery has appears to have a positive impact on the reporting, which may be attributed to an effect whereby they prompt the experimenters to consider details relating to the modality and perspective of imagery to be used.

Finally, it is important to remember that standardisation of protocols for use in motor imagery studies is essential. Indeed, as we have seen previously, in diverse ways of imagining a movement, the same neuromuscular patterns and brain areas are not activated. The lack of reporting could therefore affect the understanding, reproducibility, and translation of results found in the literature. In future experiments on motor imagery, it will therefore be important to improve the reporting of this information in future studies.

## Supporting information

Data analysis of methods in motor imagery litterature

